# Functional and biological diversity jointly shape growth and recovery of *Synechococcus* communities under stressors

**DOI:** 10.64898/2026.04.21.719904

**Authors:** Mark Holmes, Arunima Sikder, Pauline Witsel, Frederik De Laender

## Abstract

Biodiversity is expected to enhance the stability of ecological communities under environmental stress, but the relative roles of functional, interspecific, and intraspecific diversity remain poorly resolved, particularly under multiple concurrent stressors. We tested how these diversity dimensions shaped the growth and recovery of marine Synechococcus communities in a microcosm experiment manipulating strain composition across four strain-richness levels and two interspecific diversity levels under control, atrazine, warming, and combined atrazine-plus-warming treatments. Functional diversity was quantified from flow-cytometric trait data and analyzed as initial functional diversity during the stress phase and assembled functional diversity during recovery. Contrary to our expectations, higher initial functional diversity was associated with lower community growth during stress, while higher assembled functional diversity was generally associated with weaker recovery. However, these relationships depended on stressor identity and interspecific diversity: in two-species communities, the negative effects of functional diversity were reduced, and under combined stress, higher assembled functional diversity was associated with improved recovery. In contrast, intraspecific diversity consistently enhanced community growth and recovery, while interspecific diversity primarily promoted functional recovery. Together, our results show that functional, interspecific, and intraspecific diversity can influence stress responses through distinct pathways.

## 1. Introduction

The stability of biological communities is increasingly threatened by environmental stressors whose intensity and duration are exacerbated by anthropogenic global change (IPCC 2023; Pecl et al., 2017). In natural systems, however, stressors rarely act in isolation. Instead, communities are often exposed to simultaneous chemical, thermal, and biological stressors whose combined effects are difficult to predict (Nguyen et al., 2021). Such stressors can alter community structure and ecosystem functioning in ways that are difficult to predict from single-stressor responses (Jackson et al., 2021; Polazzo & Rico 2021; Orr et al., 2024). Because community stability underpins critical ecosystem functions such as primary production and nutrient cycling, identifying the mechanisms that buffer communities under multiple concurrent stressors has become a central challenge in ecology (Oliver et al., 2015).

Ecological stability is commonly characterized by two complementary components: the ability to maintain performance under stress and the ability to recover once stress is removed (Pimm 1984; Hillebrand et al. 2018). These components need not covary: communities may maintain functioning during stress but recover slowly, or conversely, decline rapidly yet recover completely (Van Meerbeek et al. 2021; Donohue 2016, 2013). Distinguishing the ecological drivers of these stability dimensions is therefore essential for predicting how communities respond to increasingly complex stress regimes.

One mechanism that may explain performance under stress acts through functional diversity (FD), defined using the distribution of functional traits within a community (Schleuter et al. 2010; Cadotte 2011; de Bello et al. 2021, Barabás et al. 2022). Communities with higher FD are often expected to better maintain ecosystem functioning under stress because trait differences can increase the likelihood that some taxa continue to perform under changing conditions (Loreau & de Mazancourt 2013). Community functioning may also depend on biodiversity dimensions other than FD. Interspecific diversity (species richness) can influence community performance through effects that are not necessarily captured by trait-space metrics alone (Tilman et al. 1997; Violle et al. 2012; Holzwart et al., 2015; MacArthur & Levins 1967; Spaak & De Laender 2021; Barabás et al. 2022). A community composed of more species, or more genotypes per species, can shape responses to stress by increasing the likelihood that some individuals persist under altered conditions (Bolnick et al. 2011; Siefert et al. 2015; Barabás et al. 2022). For example, the insurance effect occurs when taxa performing similar ecological roles respond differently to environmental change and thence improve performance (Yachi & Loreau 1999; McCann 2000; Baert et al., 2018). Similar mechanisms may operate within species at the intraspecific level and genotypic diversity may also improve stability through a similar insurance effect. This variation in responses, often termed as response diversity, can buffer ecosystem functioning when environmental conditions fluctuate because some taxa maintain performance while others decline (Elmqvist et al. 2003; Ross and Petchey 2023). Such effects may stabilize communities through demographic buffering rather than through changes in functional trait space. Interspecific and intraspecific diversity can therefore contribute to community performance through distinct mechanisms than FD, yet how these biodiversity dimensions impact recovery has not yet been jointly evaluated.

Recovery implies a potential restructuring due to historical conditions. Stress exposure may selectively eliminate trait variants and thereby compress or reorganize the functional trait space of a community (Mouillot et al. 2013; Díaz et al. 2016; Fischer et al. 2016). As a result, the FD present after stress may differ substantially from the FD present before it. This distinction is important because the FD initially present in a community may foster performance during stress, whereas FD after stress - here termed assembled FD - may hamper recovery (Baert et al. 2017, Pennekamp et al., 2018). Distinguishing between initial FD and assembled FD is therefore needed to understand how biodiversity influences both growth and recovery.

Despite increasing interest in biodiversity-stability relationships (Campbell et al., 2011, Craven et al., 2018; Eisenhauer 2024), there are currently insufficient studies that simultaneously test for effects of interspecific, intraspecific and functional diversity on stability under combined stress (Palacio et al., 2025). To address this, we experimentally manipulated species diversity and strain diversity in phytoplankton communities, specifically *Synechococcus* sp., a globally distributed primary producer in marine ecosystems, exposed to two environmental stressors in combination as well as in isolation. We quantified how intraspecific, interspecific and initial functional diversity (FD) influenced community performance and FD change during stress exposure. We then examined how intraspecific, interspecific, diversity and assembled functional diversity influenced the recovery of growth and function following stress removal.

We tested the following hypotheses. First, we hypothesized that initial FD increases community growth during stress exposure (H1). Second, we hypothesized that assembled FD improves recovery of growth and FD once stressors are removed (H2). Third, because of response diversity, we expected inter- and intraspecific diversity to have a positive effect on community performance and recovery. (H3). Finally, we expected the effects of biodiversity and functional diversity on community performance and recovery to vary with stressor identity, producing different outcomes under thermal, chemical, and combined stress (H4).

## 2. Materials and Methods

We conducted a microcosm experiment to test the effects of interspecific and intraspecific diversity and functional diversity (FD) on community responses to stress. We used four strains of marine *Synechococcus* sp. (Table 1), selected to represent two clades (hereafter “species”) and two pigmentation types (Six et al. 2007, Farrant et al. 2016, Grebert et al. 2018, Sliwinska-Wilczewska et al. 2020). All strains were obtained from the Roscoff Culture Collection. Prior to the experiment, strains were maintained under control conditions in 100 mL Erlenmeyer flasks.

**Table 1:** information on the four strains of *Synechococcus* sp. used for this study. All information was retrieved from the Roscoff Culture Collection website.

| Roscoff ID | Strain ID | Clade (Species) | Pigment type |
| --- | --- | --- | --- |
| RCC 2383 | *Synechococcus* sp. RS9909 | VIII | 1 |
| RCC 2434 | *Synechococcus* sp. RS9906 | VIII | 1 |
| RCC 2375 | *Synechococcus* sp. A15-46 | V | 2 |
| RCC 2524 | *Synechococcus* sp. BL_10 | V | 2 |

We manipulated inter- and intraspecific diversity by manipulating strain richness and composition within each strain richness level (Table 2): there were four mono-cultures (one strain, interspecific diversity of one), six duo-cultures (two strains, interspecific diversity of one to two), four tri-cultures (three strains, interspecific diversity of two), and one tetra-culture (four strains, interspecific diversity of two). Each community was subjected to one of four stressor treatments, which was replicated three times, yielding 180 experimental units total (48 monoculture, 72 duo-cultures, 48 tricultures, and 12 tetra-culture replicates).

**Table 2:** Information on the composition of communities used in this study. Strains refer to the RCC ID as indicated in Table 1.

| Strains | Species | Strain richness | Species richness |
| --- | --- | --- | --- |
| 2383, 2524 | VIII,V | 2 | 2 |
| 2434, 2524 | VIII,V | 2 | 2 |
| 2383, 2434 | VIII,VIII | 2 | 1 |
| 2383, 2434, 2524 | VIII,VIII,V | 3 | 2 |
| 2375, 2524 | V,V | 2 | 1 |
| 2375, 2383 | V,VIII | 2 | 2 |
| 2375, 2434 | V,VIII | 2 | 2 |
| 2375, 2383, 2524 | V,VIII,V | 3 | 2 |
| 2375, 2434, 2524 | V,VIII,V | 3 | 2 |
| 2375, 2383, 2434 | V,VIII,VIII | 3 | 2 |
| 2375, 2383, 2434, 2524 | V,VIII,VIII,V | 4 | 2 |

We did not manipulate FD directly but instead calculated it for each microcosm (section *Functional diversity computation*) and evaluated the effect of its initial value (initial FD) and assembled values (assembled FD) on community responses in a regression analysis (section *Data processing and statistical analysis*).

### 2.1. Treatments and microcosms

We tested four treatments: (1) control (ambient temperature 20°C, 0 mg L^−1^ atrazine), (2) atrazine (ambient temperature, 0.1 mg L^−1^ atrazine), (3) elevated temperature (+2°C above ambient, 0 mg L^−1^ atrazine), and (4) atrazine-temperature (0.1 mg L^−1^ atrazine and +2°C above ambient). The temperature increase was selected to avoid nonlinear threshold effects on growth, and the atrazine concentration was chosen to reduce intrinsic growth rates by approximately 50% (Holmes et al. 2025). Communities were cultured in 6 mL of PCRS-11 Red Sea medium (Rippka et al. 2000) in six-well plates (Sarstedt standard flat 6-well plate, 83.3920) housed in temperature-controlled incubators (Lovibond Thermostatic Cabinet). Cultures were maintained under white light (50 μmol photons m□^2^ s□^1^ at 6500 K; LedAquaristik Sky Bar) on a 12 h:12 h light:dark cycle and continuously mixed at 150 rpm using an orbital shaker (VWR mini shaker).

### 2.2. Experimental protocol and sampling

The 21-day experiment comprised two phases: a stress phase (days 0–10) and a recovery phase (days 11–20). Communities were inoculated at approximately 5000 cells/mL on day 0, with strains added in equal proportions in multi-strain communities. Every 24 h, microcosms were sampled under a laminar flow hood using a standardized procedure: (1) 57 μL of pure water was added to each well to compensate for evaporation; (2) 200 μL was removed from each well into a 96-well plate for flow cytometric analysis; (3) 200 μL of fresh medium containing the appropriate atrazine concentration was added to replace the removed volume; and (4) the collected sample was serially diluted to a density of 500–5000 cells/μL prior to measurement to be suitable for flow cytometry analysis. On day 10, the microcosms were subjected to a perturbation (dilution) and the stress treatments were reset. This consisted of setting the temperature to ambient levels (20°C) and replacing 50% (3 mL) of the medium of each microcosm with fresh medium (0 mg L^−1^ atrazine), reducing the atrazine concentration by 50% and acting as a perturbation to the communities (dilution).

### 2.3 Flow cytometry and data processing

Population density and cell traits were quantified using a Guava easyCyte 12HT flow cytometer. Five optical channels were used to distinguish live cells from non-living debris and to characterize proxies of cell traits (Rutten et al. 2005): forward scatter (cell size), side scatter (cell complexity), red-blue fluorescence (chlorophyll-a concentration), yellow-blue fluorescence (phycoerythrin concentration), and red-red fluorescence (phycocyanin concentration). Cell densities were computed after filtering doublets and debris from the raw flow cytometry data in R (v. 4.3.3) using the peacocq, cyanoFilter, flowCore, and flowDensity packages (R Core Team 2024, Olusoji et al. 2021, Ellis et al. 2024, Malek et al. 2023, Emmaneel et al. 2022).

### 2.4 Functional diversity computation

We computed FD from flow cytometric trait data following the approach of Olusoji et al. (2023) and Barabás et al. (2022). Only three of the five measured traits were used during the computation of functional diversity: cell size, chlorophyll, and phycoerythrin. These traits were the least redundant (i.e. lowest correlation with other traits) which reduced the impact of highly correlated traits on the computed FD. Additionally, this reduction of dimensionality served to reduce computational load and allowed for higher grid resolution. The multidimensional trait space was discretized into a grid (50 divisions per channel), and the kernel density of occupied trait space was estimated using the vine and dvine functions from the rvinecopulib R package (Nagler et al. 2025). Hill diversity (q = 1) was then calculated from the resulting kernel density estimate. To ensure comparability across microcosms, the grid boundaries were defined using the entire pooled dataset.

### 2.5. Data processing and statistical analysis

Four response metrics were calculated for each microcosm:

Growth of community density =

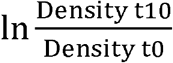

Recovery of community density =

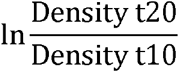

Change in functional diversity =

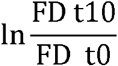

Functional recovery =

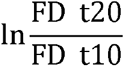

To assess how interspecific, intraspecific, and functional diversity, and stressor identity influenced community performance and FD change, we fitted four generalized least squares (GLS) models using the “nlme” package (v. 3.1-168; Pinheiro et al. 2022) in R. We used a GLS model over an ordinary least squares model to better accommodate heterogeneity in residual variance across treatment groups, modeled using a variance structure allowing residual spread to differ by treatment (varIdent, form = ~1 | treatment). All four models shared the same predictor structure: a focal FD variable (diversity at t□ or t□ □) in a three-way interaction with species diversity and stressor treatment (control, atrazine, thermal, combined), with strain diversity included as an additive covariate. The reference level for all models was set to species diversity (1), strain diversity (1), and treatment (control). Residual diagnostic plots were inspected for all models to evaluate adequacy of fit (Fig. S1).

Model coefficients and associated p-values were extracted using the “broom.mixed” package (v. 1.0.9). To characterize the direction and significance of the FD effect within each species diversity-by-treatment combination, we used the emtrends() function from the “emmeans” package (v. 1.11.2; Lenth 2023). Pairwise contrasts between slopes at the two species diversity levels were computed within each treatment using the contrast function grouped by treatment, to determine whether species diversity modulated the FD effect. This procedure was applied identically to all four models. Statistical significance was assessed at α = 0.05, with significance thresholds reported as p < 0.05 (*), p < 0.01 (**), and p < 0.001 (***). All analyses were conducted in R v. 4.5.2.

## 3. Results

### 3.1 Effects of stressors on growth and recovery of community density

Atrazine significantly reduced community growth relative to the control (β = −2.4 ± 0.1, p < 0.001, Table S1). A similar reduction was observed under the combined atrazine and thermal treatment (β = −2.4 ± 0.1, p < 0.001). Thermal stress alone had no significant effect on growth (p = 0.54).

During the recovery phase, communities previously exposed to atrazine recovered significantly better than control communities (β = 1.6 ± 0.5, p = 0.002, Table S1). Communities recovering from the combined treatment also recovered more relative to control (β = 1.3 ± 0.6, p = 0.045), whereas communities exposed to thermal stress alone did not differ significantly from the control (p = 0.13).

### 3.2 Effects of diversity on community growth

Functional diversity at the start of the 10-day stress phase (initial FD) had a significant overall negative effect on community density during the stress phase (β = −4957.6 ± 1089.1, p < 0.001, Table S1). However, this relationship depended on interspecific diversity and stressor identity. In single-species communities under control conditions, higher initial FD strongly reduced community growth. In two-species communities, this effect was reversed, resulting in a positive relationship between FD and growth (Fig 1). This difference in interspecific diversity levels was significant under atrazine (p = 0.002) and combined stress (p = 0.025), but not under control or thermal treatments.

**Figure 1:**
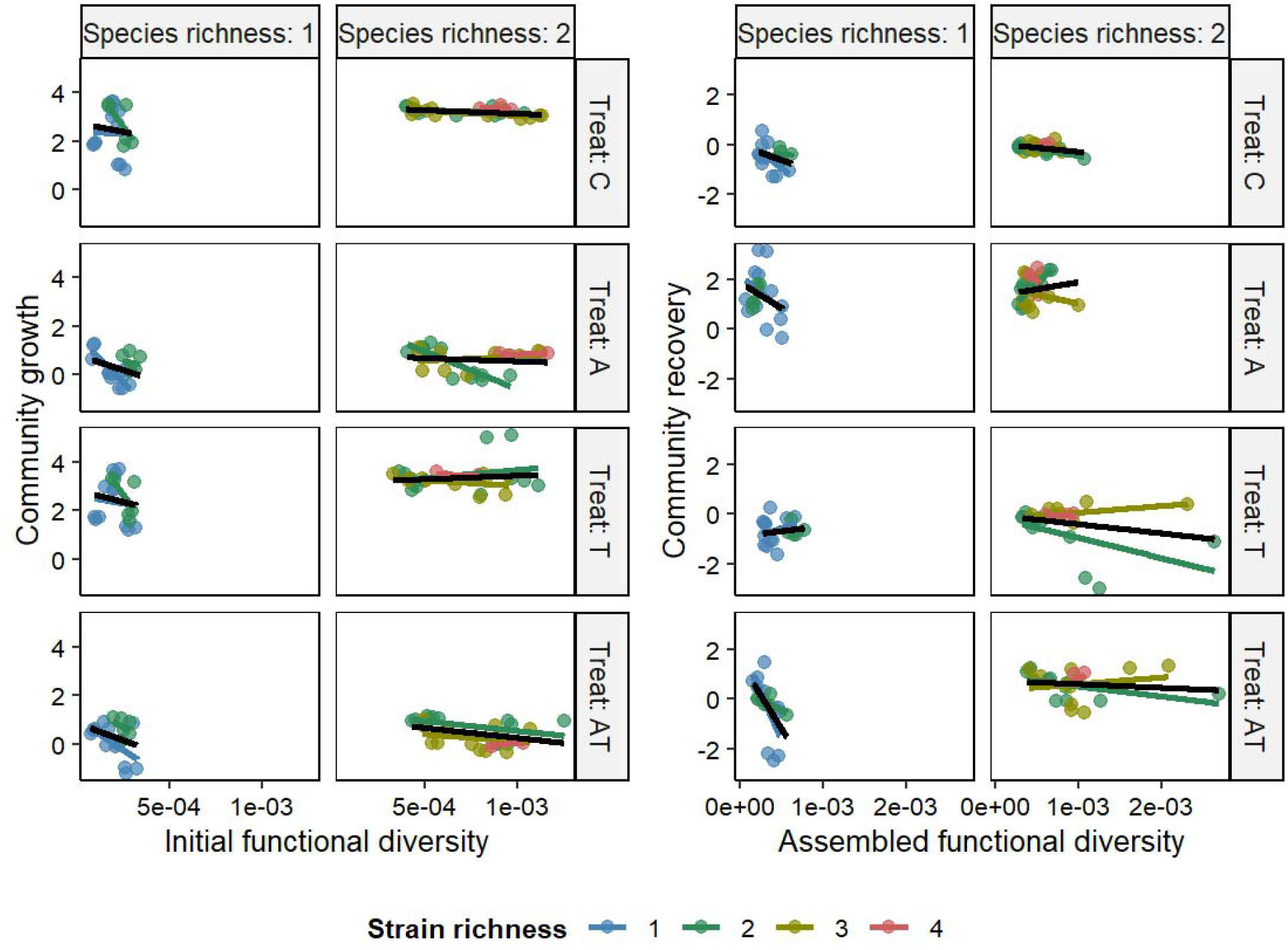
The effects of functional diversity, strain and species diversity, and stressor treatment, on community growth and recovery. Relationships are shown for exposure to control (C), atrazine (A), thermal stress (T), and combined atrazine + thermal stress (AT) treatments at two species diversity levels (1 and 2 species) and three strain diversity levels (blue = 1 strain, green = 2 strains, olive = 3 strains, and orange = 4 strains). The left panels depict how initial functional diversity relates to community growth during the stress phase, while the right panels show how assembled functional diversity (measured at the end of the stress phase) relates to density recovery after stress removal. Points represent individual microcosms and lines are fitted relationships within groups, with the black line representing the overall trend within each panel.

Interspecific diversity itself had a negative main effect on growth (β = −0.8 ± 0.4, p = 0.04, Table S1). Intraspecific diversity (the number of strains per community) positively influenced density across treatments. Duo-cultures (β = 0.8 ± 0.2, p < 0.001), tri-cultures (β = 0.7 ± 0.2, p < 0.001) and tetra-cultures (β = 0.8 ± 0.2, p < 0.001) had better growth than monocultures. These effects were consistent across treatments and did not significantly interact with stressor identity.

### 3.3 Effects of diversity on community recovery

FD after the 10-day stress phase (assembled FD) had a significant overall negative effect on community recovery (β = −1829.9 ± 711.2, p = 0.01, Table S2). (Fig 1). However, treatment type and interspecific diversity modified this trend.

Pairwise contrasts indicated that the effect of assembled FD differed significantly between single and two-species communities following exposure to the combined stressor (p = 0.003, Table S5), but not after atrazine or thermal stress alone (Fig. 3). This pattern was reflected in the significant three-way interaction in the GLS model, where two-species communities recovering from combined stress exhibited a reversal of the overall negative relationship between assembled FD and community recovery (β = 3851, p = 0.036, Table S2). Interspecific diversity alone did not significantly affect community recovery (p = 0.34, Table S2). Intraspecific diversity had a positive effect on recovery, with tri-cultures (β = 0.4 ± 0.2, p = 0.04) and tetra-cultures (β = 0.5 ± 0.2, p = 0.01) growing better, while duo-cultures did not differ significantly from monocultures.

### 3.4 Effects of stressors on functional diversity change and functional recovery

Treatments significantly influenced changes in FD during the stress phase (Table S3). Atrazine negatively influenced FD change relative to the control (β = −0.3 ± 0.1, p < 0.001). In contrast, thermal stress positively influenced FD change (β = 0.3 ± 0.1, p < 0.001), and the combined atrazine and thermal treatment also resulted in a small but significant positive effect on FD change (β = 0.2 ± 0.1, p = 0.03).

During the recovery phase, none of the treatments significantly affected functional recovery (p > 0.45, Table S4).

### 3.5 Effects of diversity on functional diversity change

Initial FD negatively affected FD change during the stress phase (Table S3). Communities with higher initial FD experienced larger declines (β = −4178.6 ± 808.5, p < 0.001). This relationship differed within interspecific diversity levels and treatments (Fig. 2). Initial FD negatively affected FD change in both single and two-species communities but the magnitude of this effect was reduced in the latter. The interaction between interspecific diversity and initial FD was positive (β = 2830.1 ± 827.0, p < 0.001), indicating that species diversity partially buffered the loss of FD in initially diverse communities.

**Figure 2:**
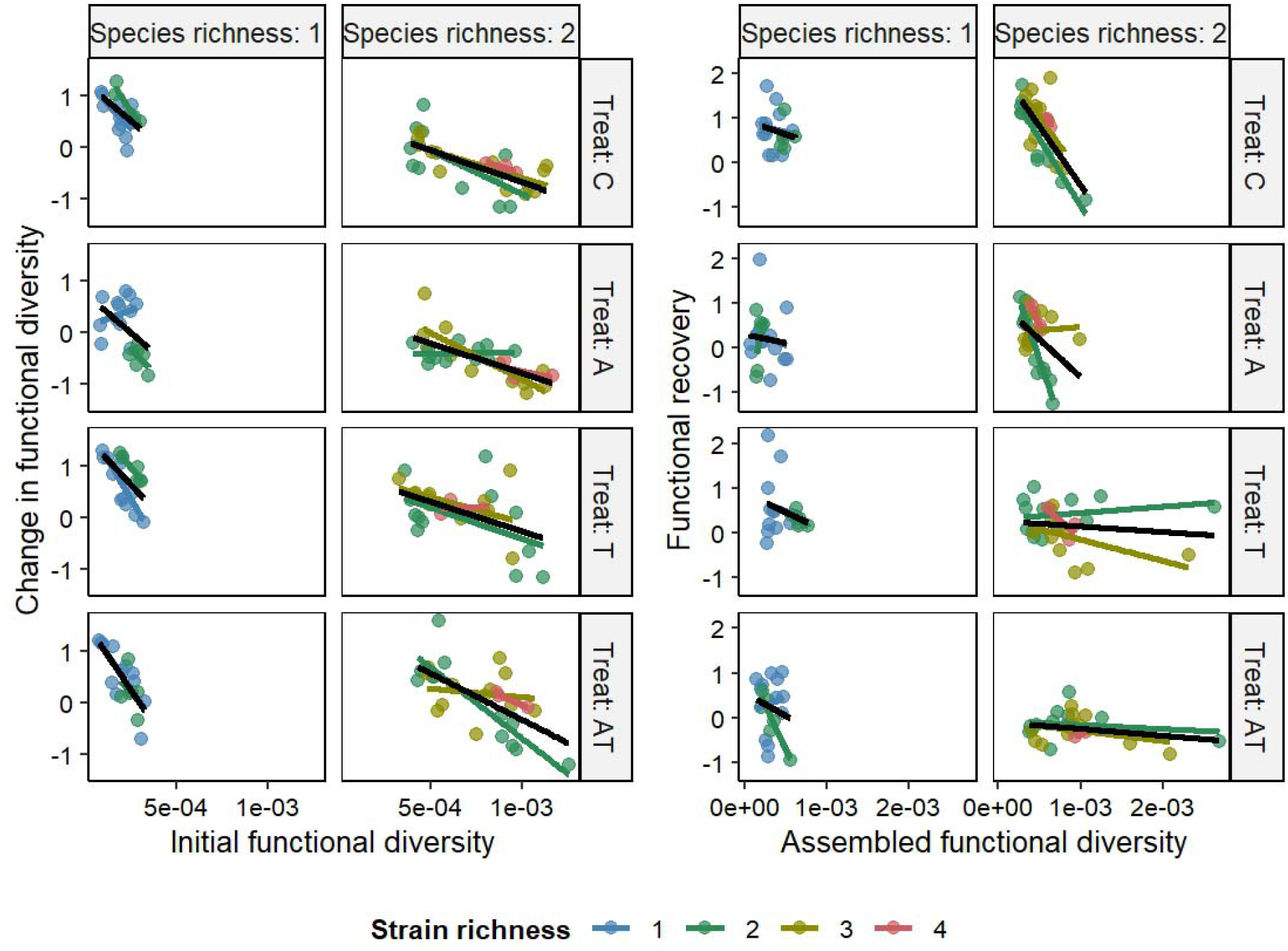
The effects of functional diversity, strain and species diversity, and stressor treatment, on change and recovery of functional diversity. Relationships are shown for exposure to control (C), atrazine (A), thermal stress (T), and combined atrazine + thermal stress (AT) treatments at two species diversity levels (1 and 2 species) and three strain diversity levels (blue = 1 strain, green = 2 strains, olive = 3 strains, and orange = 4 strains). The left panels depict how initial functional diversity relates to change of functional diversity during the stress phase, while the right panels show how assembled functional diversity (measured at the end of the stress phase) relates to recovery of functional diversity after stress removal. Points represent individual microcosms and lines are fitted relationships within groups, with the black line representing the overall trend within each panel.

Slope contrasts between interspecific diversity levels showed that this buffering effect was significant under the combined stress treatment (p = 0.022, Table S5), but not under the other treatments. Intraspecific diversity did not significantly influence FD change during the stress phase.

### 3.6 Effects of diversity on functional recovery

Assembled FD had an overall non-significant, slightly negative effect on functional recovery (β = −152.3, p = 0.88, Table S4). However, interspecific diversity significantly influenced functional recovery overall (β = 1.5 ± 0.5, p = 0.01, Table S4). Specifically, two-species communities exhibited better functional recovery than single species (Fig 3). The relationship between assembled FD and functional recovery further varied among treatments. In single-species communities, assembled FD did not significantly influence recovery under any treatment. In two-species communities, assembled FD negatively affected recovery under control conditions and following atrazine exposure. However, assembled FD did not affect functional recovery under combined and thermal stress. Under these treatments, a higher assembled FD was associated with greater functional recovery relative to control (combined: β = 3529, p = 0.02; thermal: β = 2959, p = 0.03, Table S4). Intraspecific diversity had no detectable effect on functional recovery.

**Figure 3:**
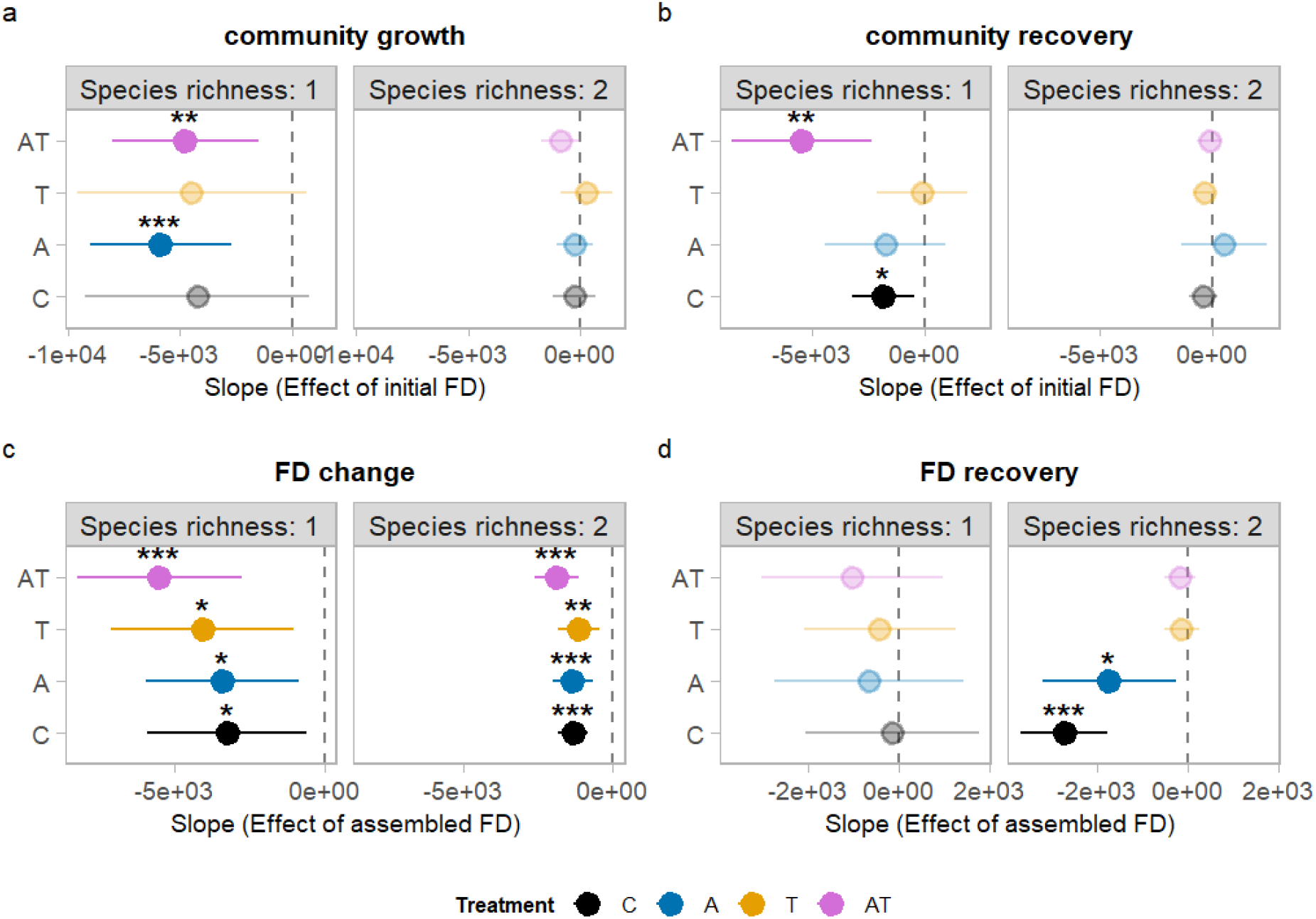
Effect of initial and assembled functional diversity (FD) on community responses across treatments and species diversity levels. Each point represents the estimated slope of the relationship between functional diversity and the response variable (community density or FD) within a given treatment and species diversity level, derived from generalized least squares models. Horizontal lines indicate 95% confidence intervals. Negative slopes indicate that higher functional diversity is associated with reduced community performance; slopes not significantly different from zero indicate no detectable FD effect. Results are shown separately for one-species (top row) and two-species communities (bottom row), and for community density and functional diversity during the stress phase (days 0–10) and recovery phase (days 11–20). Point colors denote treatment: control (C, black), atrazine (A, blue), thermal stress (T, yellow), and combined stressor (AT, pink). Asterisks indicate slopes significantly different from zero (* p < 0.05, ** p < 0.01, *** p < 0.001).

## 4. Discussion

Functional diversity (FD) influenced community responses to stress, but its effects depended strongly on stressor identity and species diversity. Contrary to our expectation that a higher initial FD would promote community growth under stress (H1), we found that initial FD reduced community growth during stress exposure. During recovery, assembled FD also had a negative effect on community growth and no effect on functional recovery reflecting a limitation on functional organization after stress removal. (H2). These negative relationships with initial and assembled FD were buffered by interspecific and intraspecific diversity consistently for both community growth and recovery (H3). Finally, as expected, these outcomes differed based on stressor identity, with stressors in isolation having a different effect than combined stressors (H4).

### 4.1 Stressor identity determines community growth and recovery

Community growth differed based on treatments, indicating that stressor identity shaped community performance under stress. Atrazine suppressed growth during the stress phase, consistent with its mode of action as a photosystem II inhibitor that constrains photosynthetic energy acquisition and phytoplankton productivity (Yang et al., 2021). In contrast, warming alone had little effect on growth relative to the control (Table S1). Temperature changes can alter phytoplankton physiology and community dynamics without necessarily imposing strong reductions in growth (Anderson et al., 2022).

Communities previously exposed to atrazine exhibited greater recovery than control communities (Table S2). One possible explanation is that chemical stress altered trait distributions in ways that changed which traits contributed most to growth after stress removal. Chemical stressors can modify response-effect relationships such that strains conferring tolerance under stress differ from those maximizing growth under control conditions (Mensens et al., 2017). More generally, trade-offs between stress tolerance and growth are widely observed across organisms (Schucht & Blasius 2025). Under this scenario, atrazine-tolerant strains may maintain limited growth during stress, whereas faster-growing strains experience stronger inhibition because photosynthesis is disrupted. Once the stressor is removed, these fast growers may rapidly resume growth and drive community recovery. Because strain identities were not tracked through time, this mechanism remains a hypothesis rather than a demonstrated process.

### 4.2 Negative effects of initial functional diversity on community growth

A higher initial functional diversity (FD) was associated with reduced community growth during stress (Table S3), suggesting that trait differentiation imposed competitive costs rather than complementarity benefits, contrary to H1. Trait differences can generate both niche differences and fitness differences, although not necessarily in conjunction (Mayfield and Levine, 2010; Spaak and De Laender 2021). Under atrazine, which constrains photosynthetic energy availability, the most functionally specialized strains were also likely the most metabolically vulnerable, contributing disproportionately to performance loss (Violle et al., 2017; Mouillot et al., 2013). Trait-based biodiversity theory similarly shows that ecosystem functioning can be shaped by opposing mechanisms, where selection driven by fitness differences can override complementarity effects (Cadotte, 2017).

This perspective explains why the negative relationship between FD and growth was strongest in single-species but weaker in two-species communities (Table S1 and S2). Within single-species communities, trait variation among strains may primarily reflect differences in competitive performance or stress tolerance, potentially increasing dominance by specific phenotypes under stress. By contrast, differences among species may be more likely to correspond to niche differentiation, allowing some of the effects of fitness differences to be offset by partial complementarity. Although our experiment cannot directly separate niche from fitness differences, the pattern is consistent with theoretical work showing that increasing diversity does not necessarily increase niche differences and may instead increase fitness differences (Spaak et al., 2021).

Intraspecific diversity consistently promoted community growth without affecting FD change and recovery (Table S1-S4). This pattern suggests that intraspecific diversity enhanced growth primarily through demographic buffering: increasing the number of strains increases the likelihood that some strains maintain growth under stress and resume growth once conditions improve.

### 4.3 Assembled functional diversity constrained community recovery

Communities with higher assembled FD at the end of the stress phase showed weaker recovery across most conditions (Table S2). This reduction in community recovery likely shows that communities retaining broader functional variation after stress had less scope for further trait reorganization and therefore showed less growth once stressors were removed (H2). Stress can reorganize communities by altering trait distributions and thereby changing which ecological strategies remain available during recovery (Mouillot et al., 2013; de Bello et al., 2021).

Under chemical stress this asymmetry was most evident: communities exposed to atrazine, though functionally impoverished after stress, contained fast-growing, tolerant strains able to rapidly recover after stressor removal, leading to greater recovery despite low assembled FD. This effect was reversed under specific conditions: two-species communities recovering from the combined stressor, higher assembled FD instead enhanced recovery (β = 3851, p = 0.036; Table S2), suggesting a possible interaction between atrazine’s filtering effect and warming’s diversifying effect produced assembled communities with both greater tolerance and greater FD than either stressor would generate independently. Intraspecific diversity continued to promote density recovery (strain diversity 4: p = 0.012, Table S2), suggesting demographic buffering as communities containing more strains are more likely to retain strains capable of sustaining growth post-stress (Bolnick et al., 2011; Siefert et al., 2015).

### 4.4 Functional diversity change during stress depended on stressor identity

FD changed during the stress phase even in control communities, indicating that community trait structure was inherently dynamic over time (Fig S2). Relative to this baseline change, atrazine shifted FD in a more negative direction whereas warming shifted it in a more positive direction (Table S3). Chemical stress may have constrained the range of physiological strategies capable of sustaining growth by limiting energy acquisition through photosynthesis (Yang et al., 2021). Thermal stress, by contrast, can modify metabolic processes and physiological traits, potentially increasing trait variation among individuals (Anderson et al., 2022; Huang et al., 2025; Armin et al., 2025). The combined treatment produced intermediate patterns of FD change (Table S3), consistent with an interaction between two opposing mechanisms that moderated the FD outcomes of either stressor acting alone (Jackson et al., 2021; Fig. 2). These contrasting outcomes demonstrate that the mode of action of a stressor, governed whether stress contracted or expanded community functional trait space.

### 4.5 Negative effect of initial functional diversity on functional diversity change during stress

Communities with higher initial FD experienced more negative FD change over time (Table S4, Fig. 2). Because FD change was also negative in control communities, this pattern reflects temporal changes in trait distributions that occur even in the absence of stress. Trait convergence through time can arise from competitive exclusion or environmental filtering that favor specific trait combinations as communities develop (Mayfield and Levine, 2010; Kraft et al., 2015). In this context, stress may modify the trajectory of this convergence rather than initiate it.

Interspecific diversity was associated with more negative FD change during the stress phase (Fig S1). One possible explanation is that communities starting with larger trait space may experience greater convergence as competitive interactions and environmental constraints favor a narrower subset of trait combinations over time. Thus, the magnitude of FD loss may scale with the amount of FD initially present. Because strain identities were not tracked through time, the specific competitive dynamics driving this convergence (whether through selective elimination of specific strains or through phenotypic plasticity within surviving strains), remain a hypothesis rather than a demonstrated process, and represent a target for future work.

### 4.6 Functional recovery primarily shaped by interspecific diversity

Interspecific diversity promoted functional recovery once stress was removed (species diversity 2: p = 0.006; Table S4), reversing its negative stress-phase effect: strains suppressed or reduced below detection thresholds during stress, may have re-established once selective pressure was relaxed, restoring functional diversity to levels unattainable by single-species communities (Elmqvist et al., 2003; Loreau et al., 2021; Loreau & de Mazancourt, 2013). The contribution of assembled FD to functional recovery was stressor-dependent and composition-specific. In two-species communities recovering from the combined stress, higher assembled FD enhanced functional recovery (β = 3529, p = 0.024; thermal: β = 2959, p = 0.030; Table S4), suggesting that the trait combinations retained under combined stress were particularly well positioned to diversify further once stressors were removed. Rather than implying a universal positive effect of assembled FD on recovery, this result indicates that recovery trajectories depend on which trait combinations remain after stress and how those combinations interact with the post-stress environment.

### 4.7 Perspectives

This study advances mechanistic understanding of diversity-stability relationships by explicitly considering how multiple diversity types interact under stress, thereby revealing the link between biodiversity, functional diversity, and community performance. Different kinds of diversity had different effects: net negative effect of both initial and assembled FD; net positive effects of inter and intraspecific diversity; and case-specific effects of stressor identity. Taken together, these findings indicate that community performance and functional responses to stress can become decoupled, with communities recovering more rapidly than they regain functional trait diversity. These findings carry direct relevance for predicting ecosystem trajectories under global change, where chemical and thermal stressors increasingly co-occur and where recovery in community performance may depend on how functional diversity is reorganized during stress. Future work should characterize intraspecific trait distributions to identify which strain-level properties confer vulnerability or resilience and extend temporal scope to determine whether limitations on functional recovery are transient or reflect enduring reorganization of community trait structure.

## Author Contributions

Mark Holmes (data curation : lead, conceptualization: equal, analysis : equal, writing: equal); Arunima Sikder (conceptualization: equal, analysis: equal, writing: lead); Pauline Witsel (data curation: equal); Frederik de Laender (conceptualization: equal, analysis: equal, writing: equal, supervision: lead).

