## Supplementary Materials for "Functional and biological diversity jointly shape growth and recovery of *Synechococcus* communities under stressors"

**Table S1**: Model output of community growth (y~ diversity * treat * species richness+ strain richness, varIdent (1|treat))

| **term** | **estimate** | **std.error** | **statistic** | **p.value** | **significance** |
| --- | --- | --- | --- | --- | --- |
| (Intercept) | 3.047685 | 0.533479 | 5.712853 | 5.23828E-08 | *** |
| diversity_0 | -4236.79 | 2565.112 | -1.6517 | 0.100545213 |  |
| species_richness2 | -0.38538 | 0.679622 | -0.56705 | 0.571468729 |  |
| treatA | -1.76223 | 0.634033 | -2.7794 | 0.006094535 | ** |
| treatAT | -2.03894 | 0.64525 | -3.15993 | 0.001885451 | ** |
| treatT | 0.097328 | 0.790839 | 0.123069 | 0.902205672 |  |
| strain_richness2 | 0.812922 | 0.162538 | 5.001436 | 1.47499E-06 | *** |
| strain_richness3 | 0.687017 | 0.199827 | 3.438052 | 0.000745887 | *** |
| strain_richness4 | 0.820791 | 0.250833 | 3.272265 | 0.001305513 | ** |
| diversity_0:species_richness2 | 3960.481 | 2608.618 | 1.518229 | 0.130917846 |  |
| diversity_0:treatA | -1640.04 | 2902.461 | -0.56505 | 0.572826367 |  |
| diversity_0:treatAT | -536.066 | 2988.826 | -0.17936 | 0.857883126 |  |
| diversity_0:treatT | -244.28 | 3579.533 | -0.06824 | 0.945676351 |  |
| species_richness2:treatA | -0.87349 | 0.792735 | -1.10187 | 0.272162599 |  |
| species_richness2:treatAT | -0.26372 | 0.819124 | -0.32196 | 0.747900956 |  |
| species_richness2:treatT | -0.32963 | 0.967443 | -0.34073 | 0.733752831 |  |
| diversity_0:species_richness2:treatA | 1679.674 | 2966.607 | 0.566194 | 0.57205046 |  |
| diversity_0:species_richness2:treatAT | -56.3721 | 3057.912 | -0.01843 | 0.985314798 |  |
| diversity_0:species_richness2:treatT | 788.0831 | 3660.361 | 0.215302 | 0.829804353 |  |

**Table S2:** Model output for community recovery (y ~ diversity * treat * species richness+ strain richness, varIdent (1|treat))

| **term** | **estimate** | **std.error** | **statistic** | **p.value** | **significance** |
| --- | --- | --- | --- | --- | --- |
| (Intercept) | 0.133416 | 0.257394 | 0.518335 | 0.60493634 |  |
| diversity_10 | -1830.14 | 692.2794 | -2.64365 | 0.009012772 | ** |
| species_richness2 | -0.381 | 0.386599 | -0.98552 | 0.325849024 |  |
| treatA | 1.587934 | 0.498111 | 3.187913 | 0.001722004 | ** |
| treatAT | 1.267512 | 0.60702 | 2.088091 | 0.038363775 | * |
| treatT | -0.83712 | 0.533367 | -1.5695 | 0.118494749 |  |
| strain_richness2 | 0.248598 | 0.15063 | 1.650388 | 0.100813008 |  |
| strain_richness3 | 0.388293 | 0.180143 | 2.155473 | 0.032610369 | * |
| strain_richness4 | 0.553641 | 0.218062 | 2.538922 | 0.012067736 | * |
| diversity_10:species_richness2 | 1386.652 | 762.6991 | 1.818085 | 0.070909852 |  |
| diversity_10:treatA | 92.81942 | 1575.229 | 0.058924 | 0.95308539 |  |
| diversity_10:treatAT | -3599.67 | 1712.032 | -2.10257 | 0.037058626 | * |
| diversity_10:treatT | 1693.508 | 1120.819 | 1.510956 | 0.13275998 |  |
| species_richness2:treatA | -0.3002 | 0.703117 | -0.42696 | 0.669981307 |  |
| species_richness2:treatAT | -0.58057 | 0.693009 | -0.83775 | 0.403414624 |  |
| species_richness2:treatT | 0.727063 | 0.60868 | 1.194491 | 0.234042707 |  |
| diversity_10:species_richness2:treatA | 855.2953 | 1873.601 | 0.456498 | 0.648647051 |  |
| diversity_10:species_richness2:treatAT | 3873.229 | 1762.548 | 2.197518 | 0.029411578 | * |
| diversity_10:species_richness2:treatT | -1631.81 | 1190.422 | -1.37079 | 0.17234938 |  |

**Table S3:** Model output for FD change (y ~ diversity * treat * species richness+ strain richness, varIdent (1|treat))

| **term** | **estimate** | **std.error** | **statistic** | **p.value** | **significance** |
| --- | --- | --- | --- | --- | --- |
| (Intercept) | 1.292001 | 0.278994 | 4.630934 | 7.46252E-06 | *** |
| diversity_0 | -3255.87 | 1357.988 | -2.39757 | 0.017649159 | * |
| species_richness2 | -0.8333 | 0.367539 | -2.26724 | 0.024705982 | * |
| treatA | -0.47653 | 0.399398 | -1.19312 | 0.234579189 |  |
| treatAT | 0.301942 | 0.416311 | 0.72528 | 0.469333022 |  |
| treatT | 0.331318 | 0.44466 | 0.745105 | 0.457294679 |  |
| strain_richness2 | 0.08506 | 0.111232 | 0.764709 | 0.445564315 |  |
| strain_richness3 | 0.229222 | 0.137356 | 1.668814 | 0.097097729 |  |
| strain_richness4 | 0.323341 | 0.172197 | 1.877739 | 0.062225382 |  |
| diversity_0:species_richness2 | 1930.134 | 1380.578 | 1.398062 | 0.16401744 |  |
| diversity_0:treatA | -153.205 | 1784.815 | -0.08584 | 0.931701963 |  |
| diversity_0:treatAT | -2254.71 | 1899.442 | -1.18704 | 0.236961919 |  |
| diversity_0:treatT | -808.29 | 2002.241 | -0.40369 | 0.68697461 |  |
| species_richness2:treatA | 0.35296 | 0.510627 | 0.691227 | 0.490418105 |  |
| species_richness2:treatAT | 0.590649 | 0.54561 | 1.082548 | 0.280627952 |  |
| species_richness2:treatT | -0.09122 | 0.544985 | -0.16738 | 0.867279529 |  |
| diversity_0:species_richness2:treatA | 145.6329 | 1830.956 | 0.079539 | 0.936702473 |  |
| diversity_0:species_richness2:treatAT | 1719.49 | 1951.681 | 0.88103 | 0.379614871 |  |
| diversity_0:species_richness2:treatT | 1011.498 | 2048.952 | 0.493666 | 0.622215048 |  |

**Table S4:** Model output for functional recovery (y ~ diversity * treat * species richness+ strain richness, varIdent (1|treat))

| **term** | **estimate** | **std.error** | **statistic** | **p.value** | **significance** |
| --- | --- | --- | --- | --- | --- |
| (Intercept) | 0.829635 | 0.383333 | 2.164267 | 0.031917438 | * |
| diversity_10 | -154.169 | 979.8131 | -0.15735 | 0.875170001 |  |
| species_richness2 | 1.487125 | 0.522408 | 2.846672 | 0.00499351 | ** |
| treatA | -0.39125 | 0.51856 | -0.7545 | 0.451650754 |  |
| treatAT | -0.19837 | 0.533637 | -0.37174 | 0.710578374 |  |
| treatT | -0.07564 | 0.539719 | -0.14015 | 0.888718211 |  |
| strain_richness2 | -0.19314 | 0.150575 | -1.28268 | 0.201446666 |  |
| strain_richness3 | -0.12641 | 0.183223 | -0.68992 | 0.491236944 |  |
| strain_richness4 | 0.017325 | 0.22378 | 0.077419 | 0.93838641 |  |
| diversity_10:species_richness2 | -2568.56 | 1097.277 | -2.34085 | 0.020465642 | * |
| diversity_10:treatA | -507.151 | 1499.254 | -0.33827 | 0.735601047 |  |
| diversity_10:treatAT | -884.782 | 1421.996 | -0.62221 | 0.534683374 |  |
| diversity_10:treatT | -276.083 | 1199.777 | -0.23011 | 0.818297204 |  |
| species_richness2:treatA | -0.73072 | 0.687249 | -1.06325 | 0.28926185 |  |
| species_richness2:treatAT | -2.04861 | 0.626907 | -3.26781 | 0.001324943 | ** |
| species_richness2:treatT | -1.82613 | 0.634052 | -2.88009 | 0.004516975 | ** |
| diversity_10:species_richness2:treatA | 1499.057 | 1746.56 | 0.858291 | 0.392007709 |  |
| diversity_10:species_richness2:treatAT | 3439.708 | 1515.605 | 2.269528 | 0.024563471 | * |
| diversity_10:species_richness2:treatT | 2871.002 | 1312.544 | 2.187356 | 0.03015858 | * |

**Table S5 :** Pairwise emmeans contrasts between species richness levels for the effect of initial and assembled functional diversity.

| **Model** | **Treat** | **Estimate** | **Se** | **Df** | **P.value** | **Significance** |
| --- | --- | --- | --- | --- | --- | --- |
| Density (stress) | Control | -3932 | 2698 | 40 | 0.153 | - |
| Density (stress) | Atrazine | -5474 | 1700 | 45 | 0.002 | ** |
| Density (stress) | Combined | -4062 | 1744 | 37 | 0.025 | * |
| Density (stress) | Thermal stress | -4777 | 2770 | 40 | 0.092 | - |
| Density (recovery) | Control | -1405 | 784 | 43 | 0.080 | - |
| Density (recovery) | Atrazine | -2273 | 1703 | 36 | 0.190 | - |
| Density (recovery) | Combined | -5256 | 1642 | 36 | 0.003 | ** |
| Density (recovery) | Thermal stress | 249 | 1082 | 44 | 0.819 | - |
| FD (stress) | Control | -1919 | 1428 | 39 | 0.187 | - |
| FD (stress) | Atrazine | -2101 | 1391 | 40 | 0.139 | - |
| FD (stress) | Combined | -3590 | 1501 | 40 | 0.022 | * |
| FD (stress) | Thermal stress | -2917 | 1645 | 38 | 0.084 | - |
| FD (recovery) | Control | 2648 | 1128 | 42 | 0.024 | * |
| FD (recovery) | Atrazine | 1055 | 1342 | 39 | 0.436 | - |
| FD (recovery) | Combined | -881 | 1064 | 37 | 0.413 | - |
| FD (recovery) | Thermal stress | -311 | 901 | 47 | 0.731 | - |

**Fig S1 : Diagnostic plots for GLS models .** Three diagnostic panels evaluate the assumptions of the GLS model. (Left) Residuals vs. Fitted values plot: normalized residuals are plotted against fitted values to assess homoscedasticity and linearity; the dashed horizontal line at zero serves as reference, with no strong systematic pattern expected under model assumptions. (Center) Normal Q–Q plot: quantiles of the normalized residuals are plotted against theoretical normal quantiles; points falling along the diagonal line indicate that residuals are approximately normally distributed. (Right) Residual distribution histogram: the frequency distribution of raw residuals across bins, providing a visual assessment of normality and symmetry around zero.

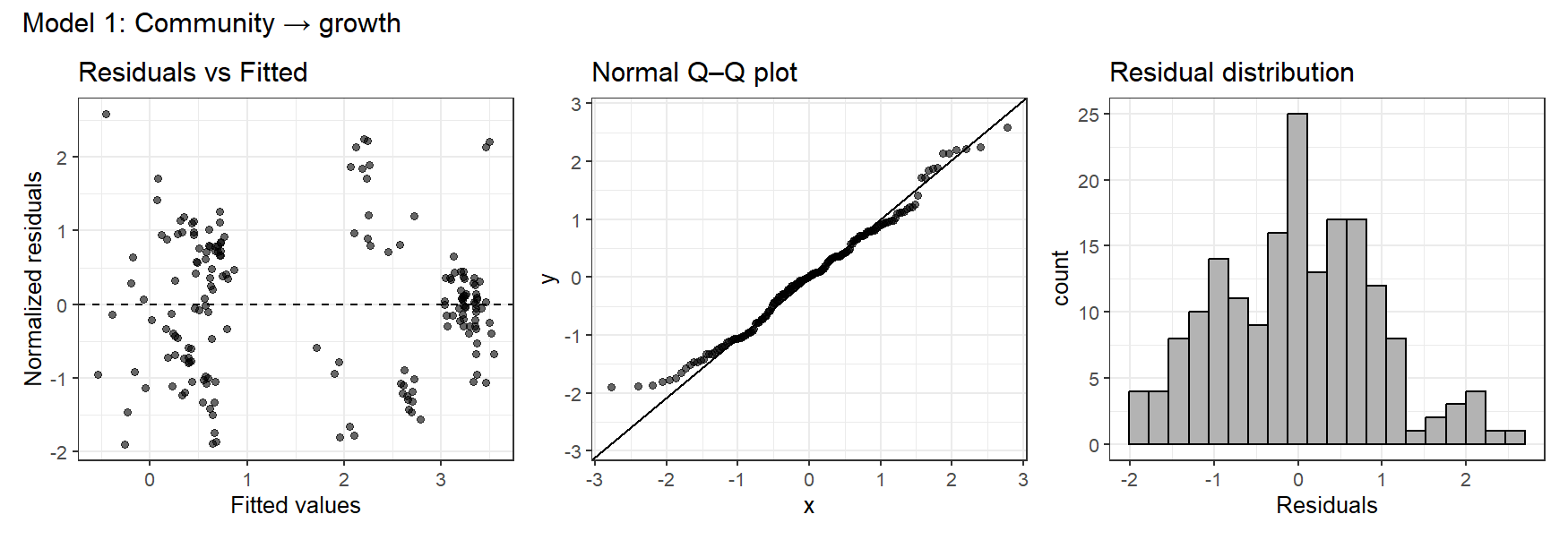

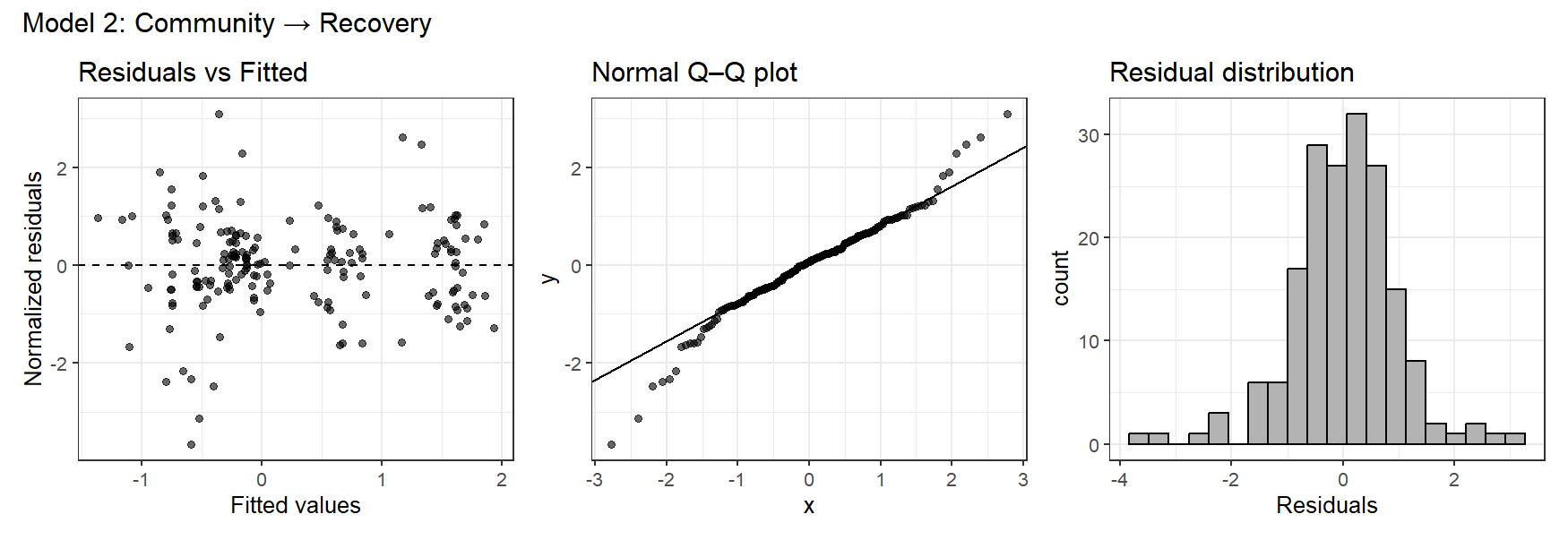

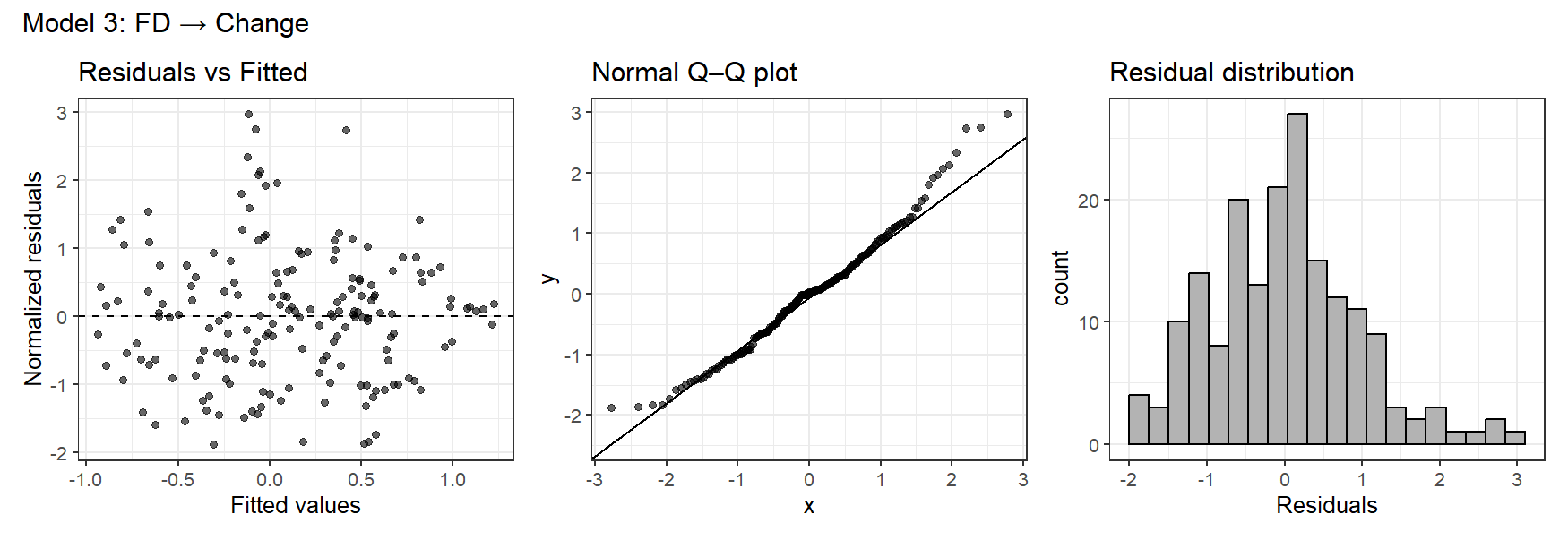

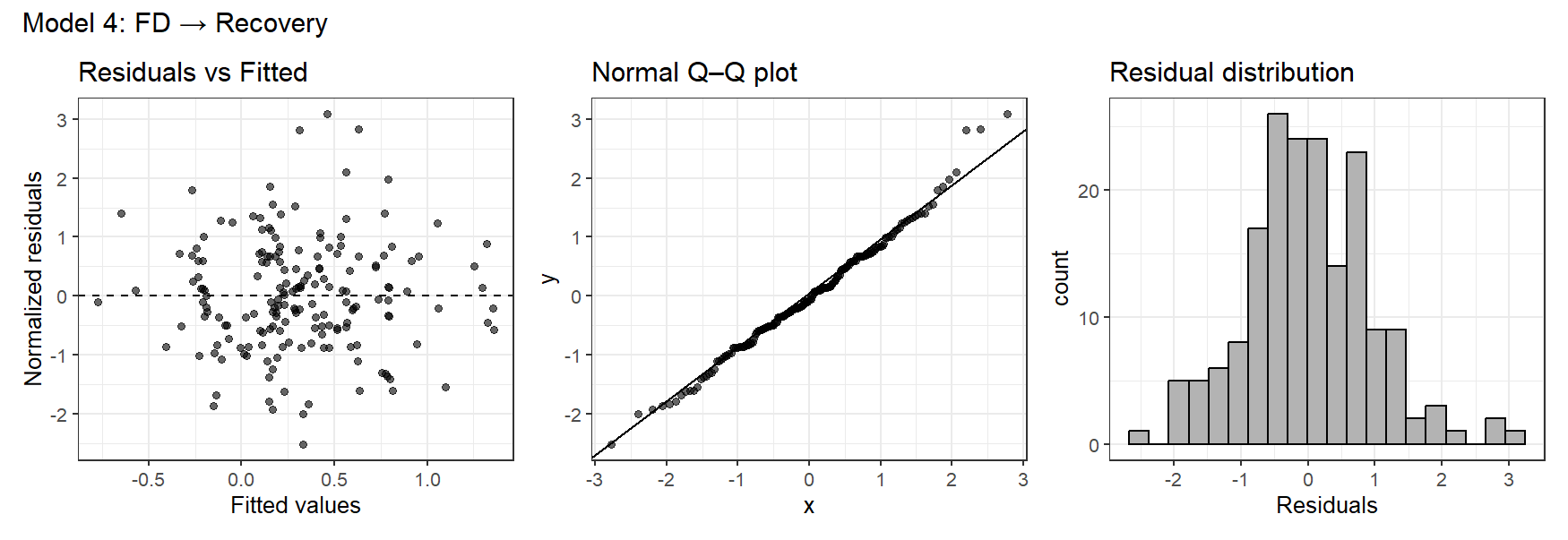

**Figure S2.** Functional diversity (FD) of *Synechococcus* sp. communities across three time points (Day 0: pre-stress, Day 10: end of stress, Day 20: end of recovery) under four stressor treatments: atrazine (A), combined atrazine and thermal stress (AT), control (C), and thermal stress (T). Boxes show the interquartile range with the median line, and whiskers extend to 1.5× the interquartile range. Individual data points are overlaid and colored by strain richness for each level of species richness.

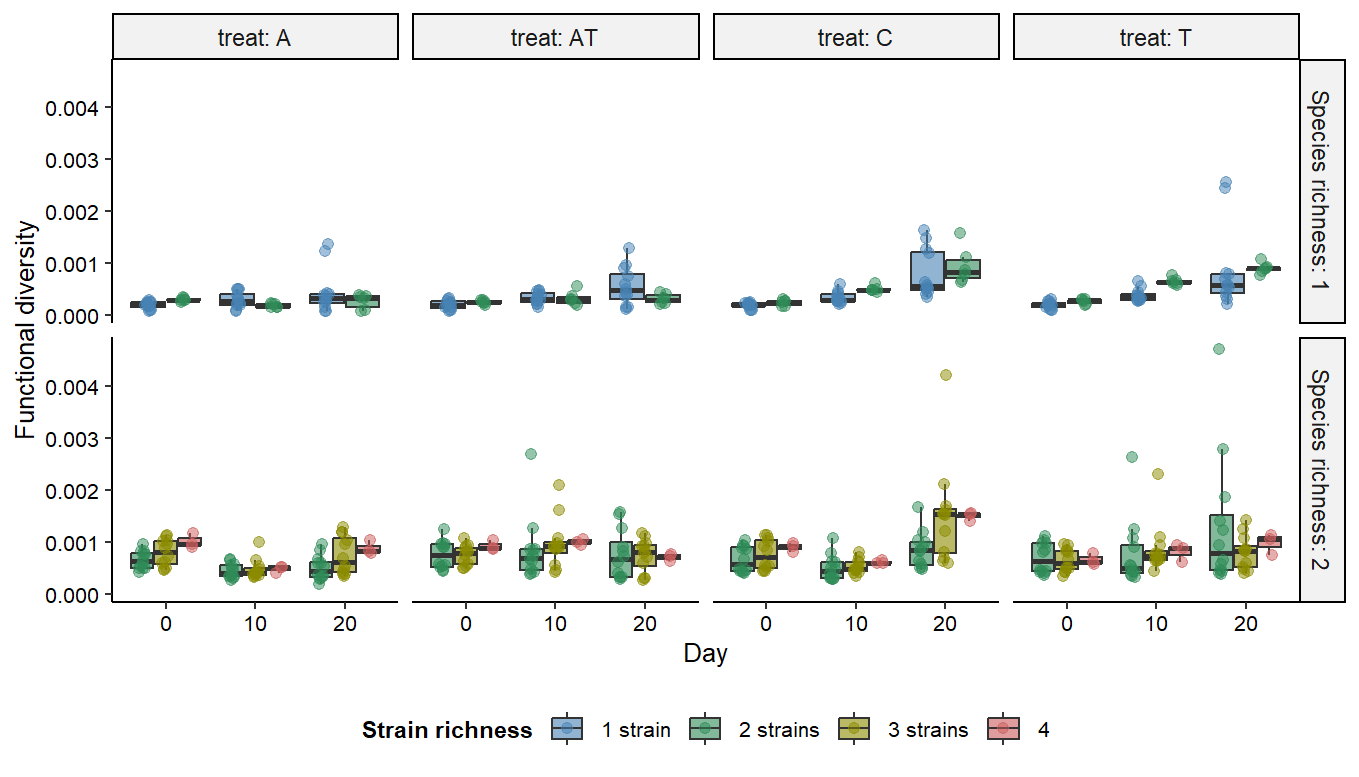
